# A looping-based model for quenching repression

**DOI:** 10.1101/085217

**Authors:** Y. Pollak, S. Goldberg, R. Amit

## Abstract

We model the regulatory role of proteins bound to looped DNA using a simulation in which dsDNA is represented as a self-avoiding chain, and proteins as spherical protrusions. We simulate long self-avoiding chains using a sequential importance sampling Monte-Carlo algorithm, and compute the probabilities for chain looping with and without a protrusion. We find that a protrusion near one of the chain’s termini reduces the probability of looping, even for chains much longer than the protrusion–chain-terminus distance. This effect increases with protrusion size, and decreases with protrusion-terminus distance. The reduced probability of looping can be explained via an eclipse-like model, which provides a novel inhibitory mechanism. We test the eclipse model on two possible transcription-factor occupancy states of the *eve* 3/7 enhancer, and show that it provides a possible explanation for the experimentally-observed *eve* stripe 3 and 7 expression patterns.

The authors declare no conflict of interests

## INTRODUCTION

Polymer looping is a phenomenon that is critical for the understanding of many chemical and biological processes. In particular, DNA looping has been implicated in transcriptional regulation across many organisms, and as a result plays a crucial role in how organisms develop and respond to their environments. While DNA looping has been studied extensively over the last several decades both experimentally (1–5) and theoretically (6–11), many aspects of looping-based transcriptional regulation remain poorly understood.

*In vivo*, the simplest looping-based regulatory architecture is comprised of a protein that interacts with a distal site via DNA looping. Such simple architectures can be found in bacteria, where, for example, σ^54^ (σ^N^) promoters are activated via such a mechanism (12–14). In eukaryotes, DNA-looping-based regulation is associated with the interaction between the core promoter and distal regulatory regions called enhancers. These ~500 bp enhancers typically contain clusters of transcription factor (TF) binding sites, and may be located from 1 Kbp to several Mbps away from their regulated promoters. Detailed studies of enhancers from several organisms (15) have revealed that TFs can upregulate, inhibit, or both upregulate and inhibit gene expression via a variety of mechanisms. For example, repressors like *D. melanogaster* Giant, Knirps, Krüppel, and Snail inhibit expression either by partial overlap of their binding sites with that of an activator, or via a short-range repression mechanism termed “quenching”, whereby TFs positioned several tens of bps either upstream or downstream from the nearest activator inhibit gene expression (16–19).

In quenching, a bound protein inhibits gene expression, apparently without any direct interaction between the protein and either the promoter or the nearest-bound activator. In the prevailing model, quenching is assumed to be a result of histone deacetylation, which is facilitated by the formation of a ~450 kDa DNA-bound chromatin remodelling complex made of a DNA binding protein such as Knirps, C-terminus binding protein (CtBP) (20–22), and the histone deacetylases (HDACs) Rpd3 (23) and Sin3 (24). However, this model falls short of providing a full description of quenching. In particular, short-range repressors such as Knirps have been shown to retain a regulatory function even without the CtBP binding domain (22, 25, 26), and tests for HDAC activity using either Rpd3 knock-outs (27) or the HDAC inhibitor trichostatin A failed to alleviate the observed repression effects (26). Consequently, the mechanistic underpinnings of quenching are still poorly understood.

In this report, we show that DNA looping can provide a mechanistic model for quenching repression. We use a modified worm-like chain model that takes excluded-volume considerations into account (28–30) to show that volume effects can substantially affect loop formation for chains that are arbitrarily long. Using this approach, we show that the excluded volumes of DNA and a protein bound within ~1 Kuhn length (~300 bp) either upstream or downstream of one of the loop termini can block the “line-of-sight” of the other terminus, generating an eclipse-like effect, which leads to a reduction in the probability of looping.

## THEORY

### DNA in the absence of bound proteins

We model the DNA as a discrete semi-flexible chain made of individual links of length *l*. A chain is described by the locations r_*i*_ of its link ends, and a local coordinate system defined by three orthonormal vectors 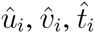 at each link, where 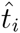 points along the direction of the ith link. We use the following notations for a specific chain configuration: *θ_i_*, _*i*_ are the zenith and azimuthal angles of 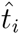 in local spherical coordinates of link *i*–1, respectively. 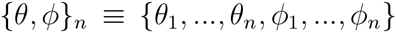 denotes all the angles until link *n*. Joint *i* is the end-point of link *i* and joint 0 is the beginning terminus of the chain. *w* is the effective cross-section of the polymer. Each chain joint is engulfed by a “hard-wall” spherical shell of diameter *w*. The total elastic energy associated with the polymer chain can be written as follows (28):

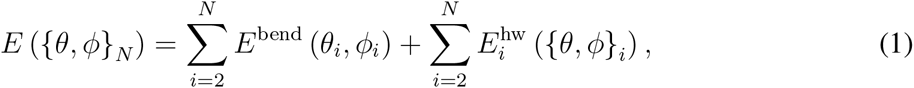

where the elastic contribution to the energy is given by:

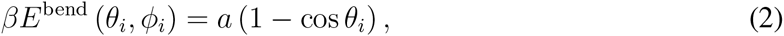

*a* is the bending constant of the polymer chain, and we have assumed azimuthal symmetry. The hard-wall contribution is given by:

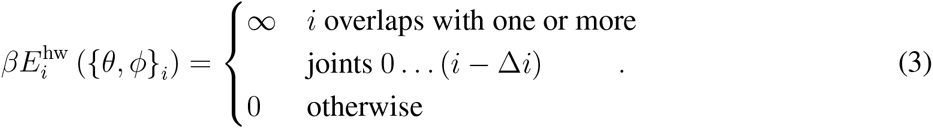

Here β = (*k*_*b*_*T*) ^1^, *k*_*b*_ is the Boltzmann factor and *T* is the temperature. In case *l* ≥ *w*, Δ*i* = 1. In case *l* < *w*, two or more consecutive spheres overlap and Δ*i* ensures that links *j* and *k* interact only if 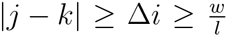. For simplicity, we disregard the twist degree of freedom in this work (see discussion in Supplementary Information Section 1.1).

### DNA in the presence of bound proteins

We model the bound proteins as hard-wall spherical protrusions positioned adjacent to the polymer chain, with radius *R*_o_ representative of the protein’s volume (Fig. 1). Since we neglect torsion effects in our present model, there is no intrinsic rotation of 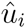 around the polymer axis. Thus, we define the orientation of the bound proteins by rotating 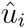 around 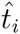. Therefore, the center of a protrusion bound to chain link *k* and rotated around the chain axis by an angle of γ_*k*_ is given by:

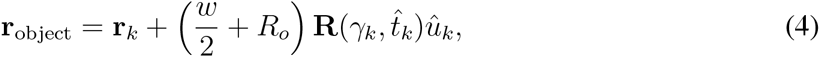

where 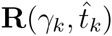 is a rotation matrix by an angle γ_*k*_ around 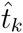 (see Fig. 1). Addition of a protrusion at link *k* slightly alters Eq. 3), requiring to test whether joint *i* overlaps with one or more joints 0 … (*i*–Δ*i*) and with the protrusion at *r*_object_, if *i* < *k*.

### Definition of looping probability ratio

We consider a chain looped if r_*N*_ is confined to a volume *δ*r around r_0_ (see Fig. 1), defined by:

1. 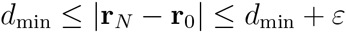.
2. 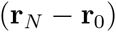 is collinear with 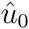 within *δω*′

For a specific choice of *δ*r, we define the probability of a polymer chain of length *L* to form a loop as (28):

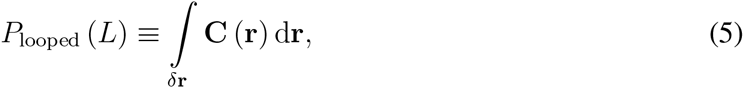

where C(r) is the probability density function of the end-to-end vector r ≡ r_*N*_ r_0_. In this work we study the effect of a bound object on the probability of the polymer to form a loop, with the looping criteria defined above. We quantify this effect by the looping probability ratio:

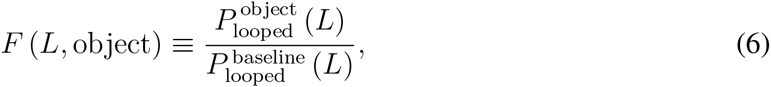

where 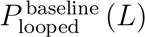 is the looping probability of the bare polymer chain and 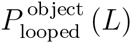 is the looping probability of the polymer chain with a protrusion bound to it.

**Figure 1:**
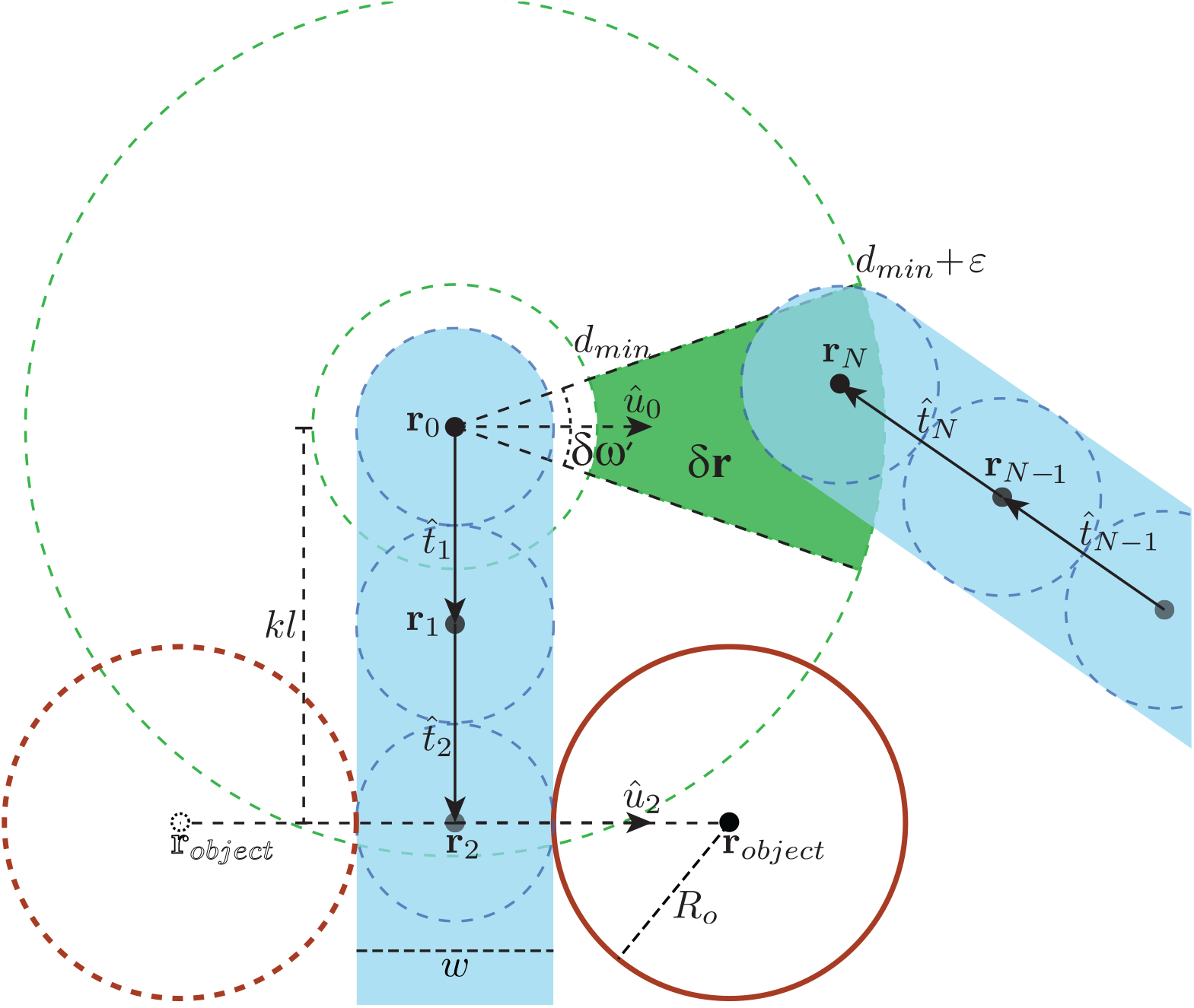
Loop and protrusion geometry. The chain (shaded in cyan) is modeled by spheres (blue dashed circles), here with link length *l* equal to diameter *w*. Looping volume δr is represented by a green wedge. The protrusion is positioned on the second link (*k* = 2). In-phase-with-δr (γ_2_ = 0°) and out-of-phase (γ_2_ = 180°) positions are illustrated by the solid and dashed red circles, respectively.

## MATERIALS AND METHODS

Linear polymers have been studied via simulations using a variety of methods (31, 32). To model DNA fully it is necessary to take into account the experimentally observed non-entangled chromosomal DNA structure (33), and to include details regarding bound proteins. Recently, polymer rings have been studied as a model system for topologically constrained polymer melts, such as chromosomal DNA (34, 35). In this work, we address looping of a linear polymer, neglecting the additional topological constraints, but including bound protrusions. We chose to simulate a linear DNA chain with a bound protein using a sequential importance sampling Monte-Carlo approach that we used previously to simulate the configurational space of bare DNA (28). To adapt our algorithm to the case of protein-bound DNA, we take into account not only the growing chain but also the location of the protrusion (see Eq. (4)). During chain generation, upon reaching link k, the simulation adds a hard-wall spherical protrusion with radius R_o_ at the location r_object_. If the protrusion overlaps any of the previously-generated chain links or protrusions, the chain is discarded. After generating the configurational ensemble, we identify the subset of “looped” chains. We provide the essential details of the simulation in this Section. Additional details can be found in the extended Materials and Methods online.

### Sequential importance sampling

Our Monte-Carlo algorithm is an off-lattice sequential importance sampling algorithm adapted from a method developed by Rosenbluth and Rosenbluth (36) to generate self-avoiding walks on a lattice (as described in (28)). We generate faithful statistical ensembles consisting of *N*_*c*_ ≈ 10^9^ self-avoiding DNA chains with bound proteins modeled as hard-wall spheres. A chain *j* in the ensemble is assigned with a Rosenbluth factor *w*_*j*_ in order to negate the biasing effect introduced by sampling of self-avoiding chains. The partition function of the ensemble takes the form of

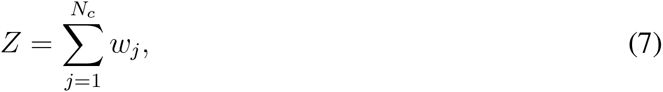

and any physical observable can then be computed from the generated ensemble by

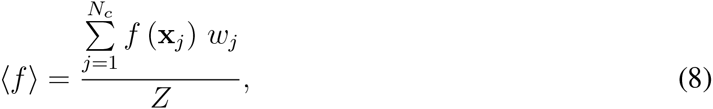

where x_*j*_ are the generalized coordinates of configuration *j*. The probability of looping is computed by

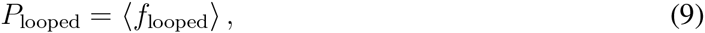

where

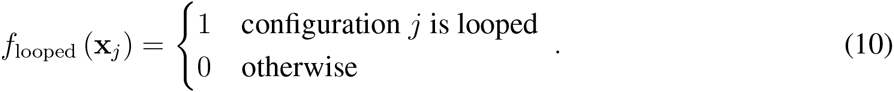

### Simulation parameters

In our simulations, *d*_min_ = *w*, ε = 2*w* and *δw*′ = 2π×0.1, unless stated otherwise. Changing these parameters did not alter the results significantly, and the relatively large ε chosen minimized noise. We simulated the DNA chain with diameter *w* = 4.6 nm and Kuhn length (7) *b* = 106 nm, where *a* is computed using (30):

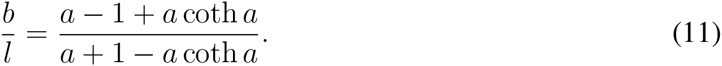

Chains were simulated in two stages as a compromise between resolution and running time. The first *N*_1_ links of the chain were simulated with link length *l*_1_ = 0.34 nm, corresponding to the length of a base-pair in dsDNA. We used 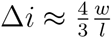 for this stage (37). All bound objects were positioned at links *k* < *N*_1_. The remaining *N*_2_ links of the chain were simulated with link length *l*_2_ = *w*. We denote the overall length of the chain by *L* = *N*_1_*l*_1_ + *N*_2_*l*_2_, and the distance along the chain of an object binding link *k* from the chain origin by *K* = *kl*_1_. For further details see Extended Materials and Methods.

### Natural system simulation

We modeled the structure of the *eve 3/7* enhancer in *D. melanogaster* based on (38). The dStat (86 kDa), Zld (146 kDa), bare Knirps (46 kDa), Knirps bound to CtBP dimers (130 kDa), and a full putative 450 kDa complex as reported by (23) were modeled as hard-wall spheres with sizes corresponding to globular proteins with radii of 3.04 nm, 3.75 nm, 2.38 nm, 3.59 nm and 5.83 nm, respectively, based on the work by (39). We used 2π / 10.5 as the DNA native twist. The chain was assumed to be rigid with respect to the twist degree of freedom. The RNA polymerase-and-cofactors complex was not modeled. Instead, for a loop to form, the chain terminus distant from the enhancer was required to be in close proximity to one of the three activators. A chain was considered looped with respect to a specific activator if it fulfilled the following conditions:

1. 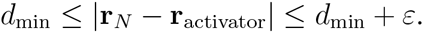.
2. 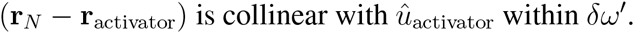.

Here r_activator_ is the center of the activator sphere (r_object_ in Fig. 1) and 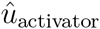 is the vector point-ing from the chain axis to the center of the activator (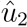 in Fig. 1). We simulated DNA chains of length 3900 bp, which corresponds to the native *eve* 3/7 enhancer geometry and its separation from the *eve* promoter. The total probability of looping was computed as the sum of the looping probabilities with respect to the individual activators. In this simulation we used *d*_min_ = *R*_chain_ + *R*_activator_ and ε = 3 nm. The simulation was relatively insensitive to ε. For *δw*′, we chose several values: *δw*′ = 2π˟0.1 corresponding to a narrow cone directed away from the chain, *δw*′ = 2π corre-sponding to a full spherical shell around the activator sphere, and *δw*′ = −2π×0.5 corresponding to a relatively wide cone directed towards the chain.

### Computation of *F*_∞_ and its error estimation

See Extended Materials and Methods.

## RESULTS

### Long-range down-regulatory effect

To model the quenching effect of a bound repressor on DNA looping, we generated configurational ensembles for DNA with a spherical protrusion of size *R*_o_ = 9.2 nm or *R*_o_ = 18.4 nm located a distance of *K* = 95 bp or *K* = 135 bp from the chain origin along the chain, oriented either in the same direction as the looping volume *δ*r or 180° from it see (Fig. 1). We plot *F* (*L*) for the various configurations of *R*_o_ and *K* in Fig. 2A. The data show that in the elastic regime (*L* ≤ *b*), protrusions bound in-phase with *δ*r (solid lines, ↑) strongly reduce the looping probability relative to that of the bare DNA, while protrusions positioned out-of-phase to *δ*r (dashed lines, ↓) increase the looping probability, as we showed previously (37). However, in the entropic regime (*L* ≫ *b*), all chains converge to values of 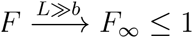. For protrusions that are bound in-phase (i.e. γ_*k*_ = 0), *F*_∞_ is distinctly smaller than one, and strongly depends on both the distance to the nearest terminus and the size of the protrusion. Conversely, for protrusions bound out-of-phase (*γ_k_* = 180°), *F*_1_ is only slightly smaller than 1, with weak dependence on both protrusion size and position.

**Figure 2:**
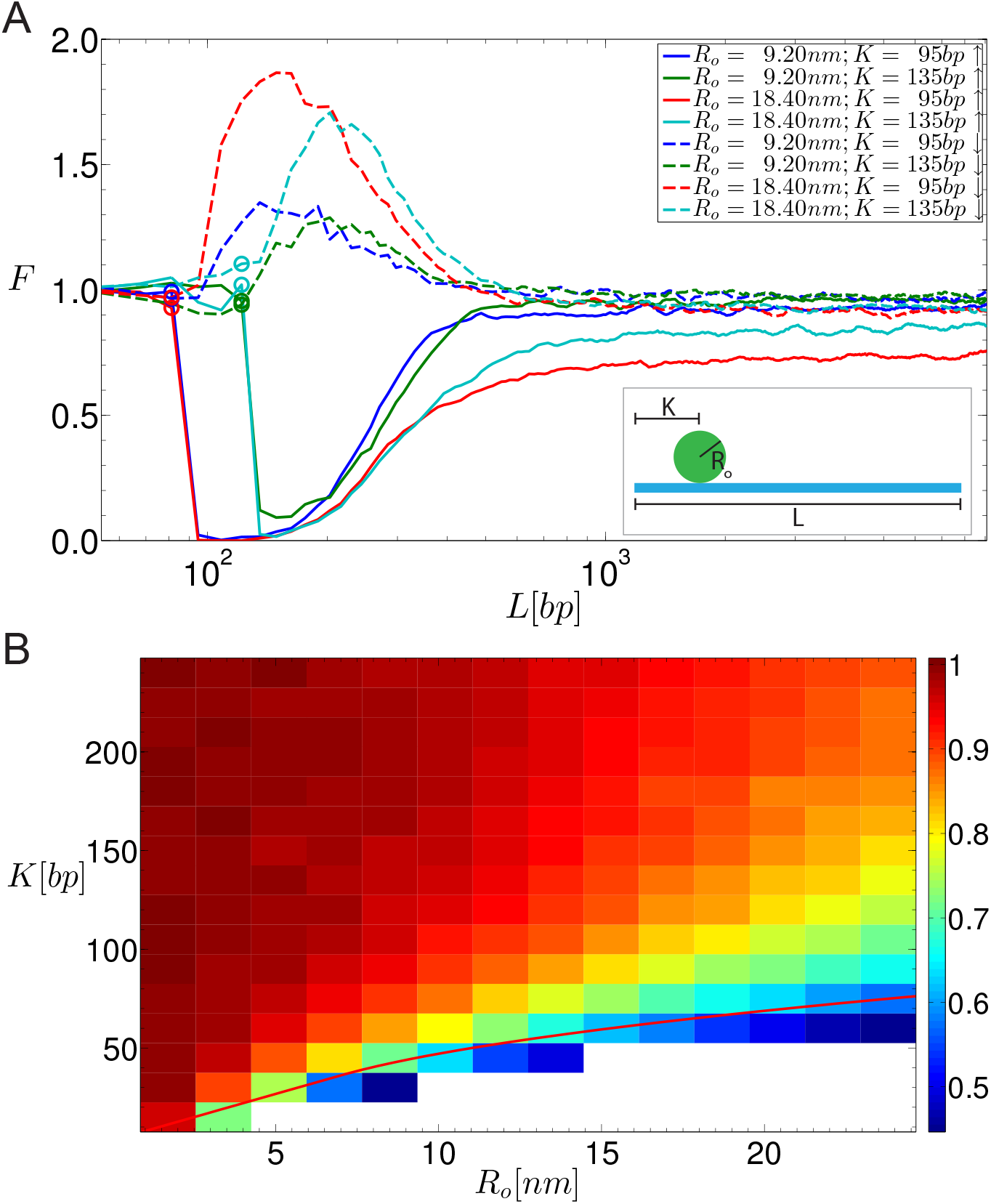
Simulating looping probability ratio *F*. A) *F* plotted as a function of chain length *L* for several values of *R*_o_ and *K*, and γ_*k*_ = 0° (solid lines) or 180° (dashed lines). The locations of the protrusions are denoted by circles on the corresponding curves. B) *F*_∞_ plotted as a function of *R*_o_ and *K*, for γ_*k*_ = 0. *F*_∞_ is calculated numerically as the average of *F* (*L*) over the range of *L* values where *F* (*L*) ≈ const. The solid red curve is a visual aid: if the segment of the chain between the origin and the protrusion location was straight, points on the red curve would result in the protrusion touching the looping volume r.

To further explore the extent of the quenching effect in the entropic or long-chain-length regime, we plot in Fig. 2B the value of *F*_∞_ as a function of a wide-range of *R*_o_ and *K*, for γ_*k*_ = 0. The heatmap shows both a non-linear decrease in *F*_∞_ as a function of protrusion size and a non-linear increase as a function of protrusion distance from the chain origin. The red line in the figure demarcates the closest possible location at which a protrusion can be bound without physically penetrating part of the looping volume *δ*r. Thus, for volumes and protrusion positions that fall below this line, a second excluded volume effect contributes to the reduction in the probability of looping, leading to a sharp increase in the overall effect. Together, the panels in Fig. 2 show that a sufficiently-large protrusion can substantially reduce the probability of looping, independent of loop length, provided that its binding site is within a small distance from either of the loop termini.

**Figure 3:**
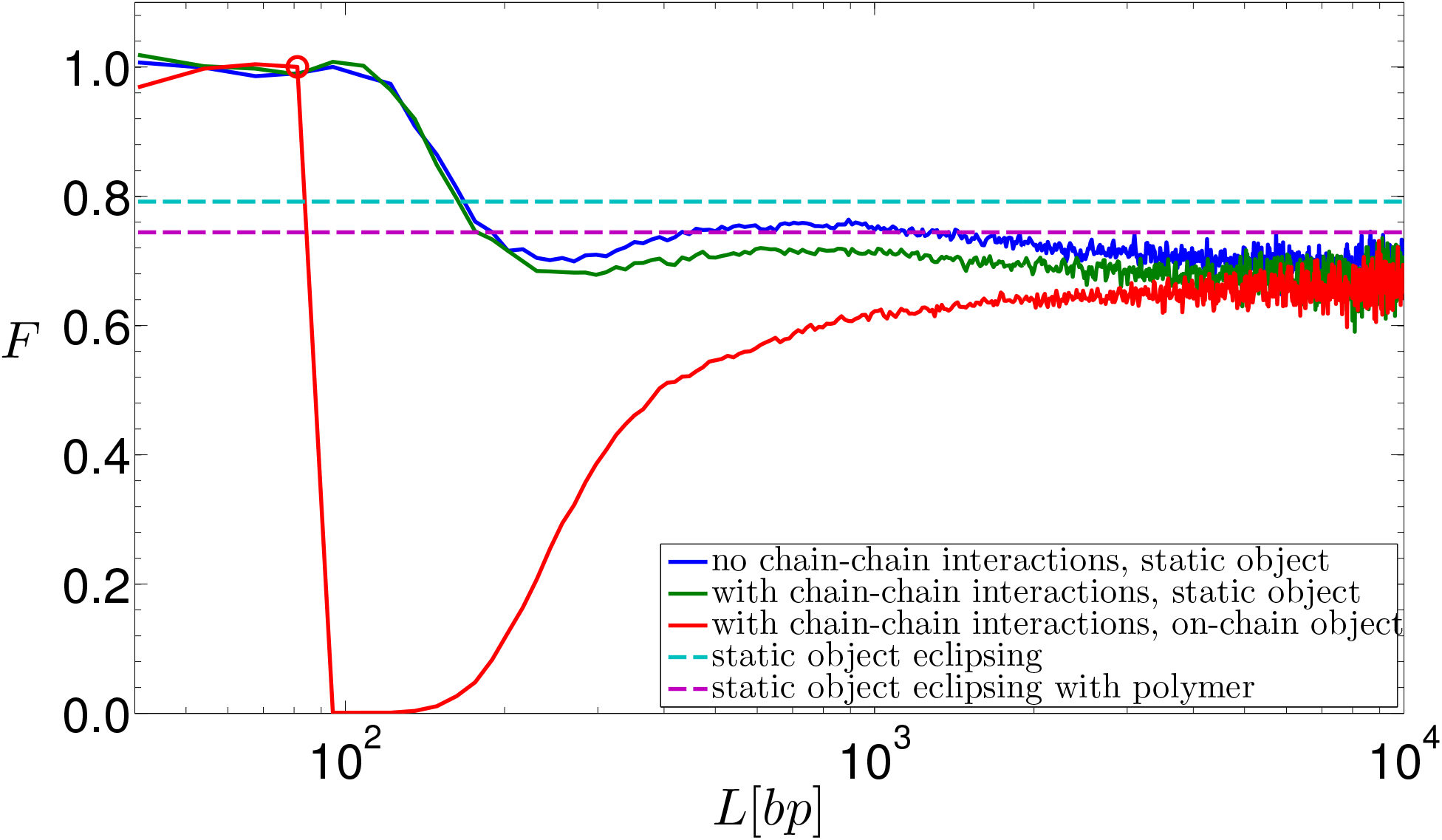
Simplified eclipse models. Simulation results for a chain without chain-chain interactions and a static object (solid blue line) are compared to estimates for *F*_∞_ of the “rod” (dashed cyan line) and “terminating-segments” (dashed magenta line) models, also for a chain without chain-chain interactions. The static object is located at 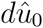, where *d* = 41.45 nm, and *R*_o_ = 23 nm. For comparison, we plot simulation results for a chain with chain-chain interactions and the same static object parameters (solid green line), and a chain with chain-chain interaction and an on-chain object (solid read line). The on-chain object is located at *K* = 94 bp (denoted by a circle).

### “Eclipsing” approximation

To understand the long-range, length-independent effect shown in Fig. 2, we examine the chains’ terminating segments of length *T* ≪ *L*. If 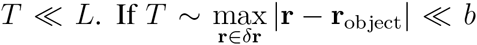, these segments resemble stiff rods, and the object obstructs the line-of-sight of one chain terminus from the other. This eclipse-like phenomenon is manifested by a reduction in the number of polymer chains that are able to reach *δ*r. This, in turn, results in a smaller P_looped_ as compared with the case in which no protrusion is present. In the entropic regime and in the absence of protrusions, the generated “rods” approach the looping volume *δ*r from all directions that are unobscured by the volume of the polymer in a homogeneous fashion (7). Due to this isotropy in the distribution of the chain termini orientations within *δ*r, the reduction in P_looped_ can be approximated by the solid angle that the eclipsing object subtends at *δ*r. Consequently, *F* (Eq. (6)) can be approximated for this “rod model” by:

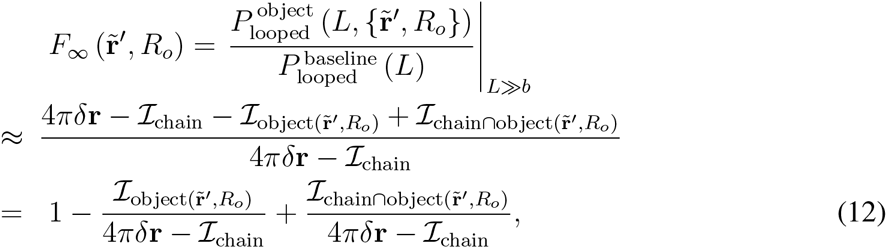

where *R*_o_ is the radius of the spherical object, 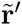 is the location of the object, which could be located statically at point r^′^ (in which case 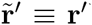), or located on the chain a distance *K* from the chain origin (in which case we use the terminology 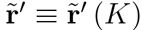 (K) figuratively to specify the progression of the protrusion along the chain). 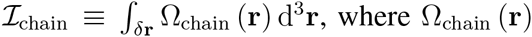 is the solidangle subtended at r by the polymer chain links.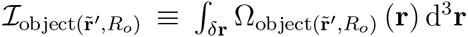, where 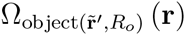 is the solid angle subtended at r by the object, and 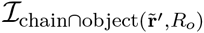 corresponds to the solid angle contained in both ℐ_chain_ and ℐ_object_.

In order to test the eclipsing hypothesis, we first computed *F*_∞_ for the case of an object statically positioned at an off-chain location 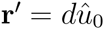, and without chain-chain interactions. In this simplified case, *F*_∞_ in Eq. (12) can be approximated by the following eclipsing expression:

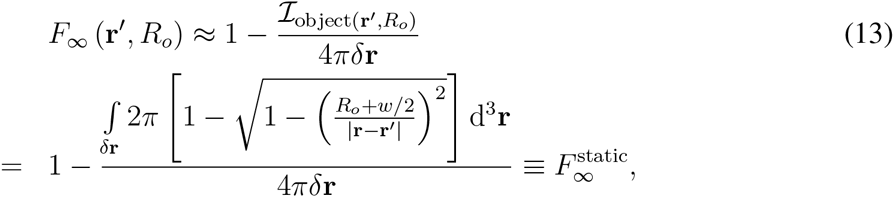

where we substituted

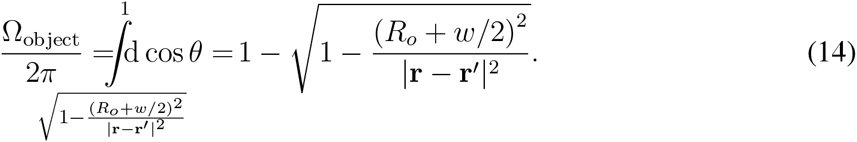

In Fig. 3, we compare the value computed from Eq. (13) (dashed cyan line) to *F* (*L*) computed by our sequential importance sampling algorithm for the same conditions (solid blue line). The data show that the eclipsing approximation 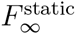 overestimates *F*_∞_. We reasoned that the main cause for this estimation error is that Eq. (13) disregards the flexible polymer nature of the chain. We ran an additional Monte-Carlo simulation to quantify the correction resulting from polymer flexibility. Here, we generated pairs consisting of an end-terminus point in δr and a direction vector of the terminal link, both distributed uniformly. Short polymer chains of length *T* originating at the chosen points were grown with their first links oriented in the chosen directions. These chains can be thought of as the terminating segments of long chains that have a uniform distribution of their end-termini in δr. We found that the probability of a flexible-polymer chain to overlap the object increased relative to the probability within the “rod model”, resulting in a decrease in the probability of the chain to form a loop (magenta dashed line in Fig. 3). Using this “terminating-segments” correction, the discrepancy between *F*_∞_ from the simulation and 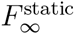 from Eq. 13 is partially accounted for. We attribute the additional reduction in the simulated *F*_∞_ to interactions between the object and the remaining *L* – *T* length of the chain.

**Figure 4:**
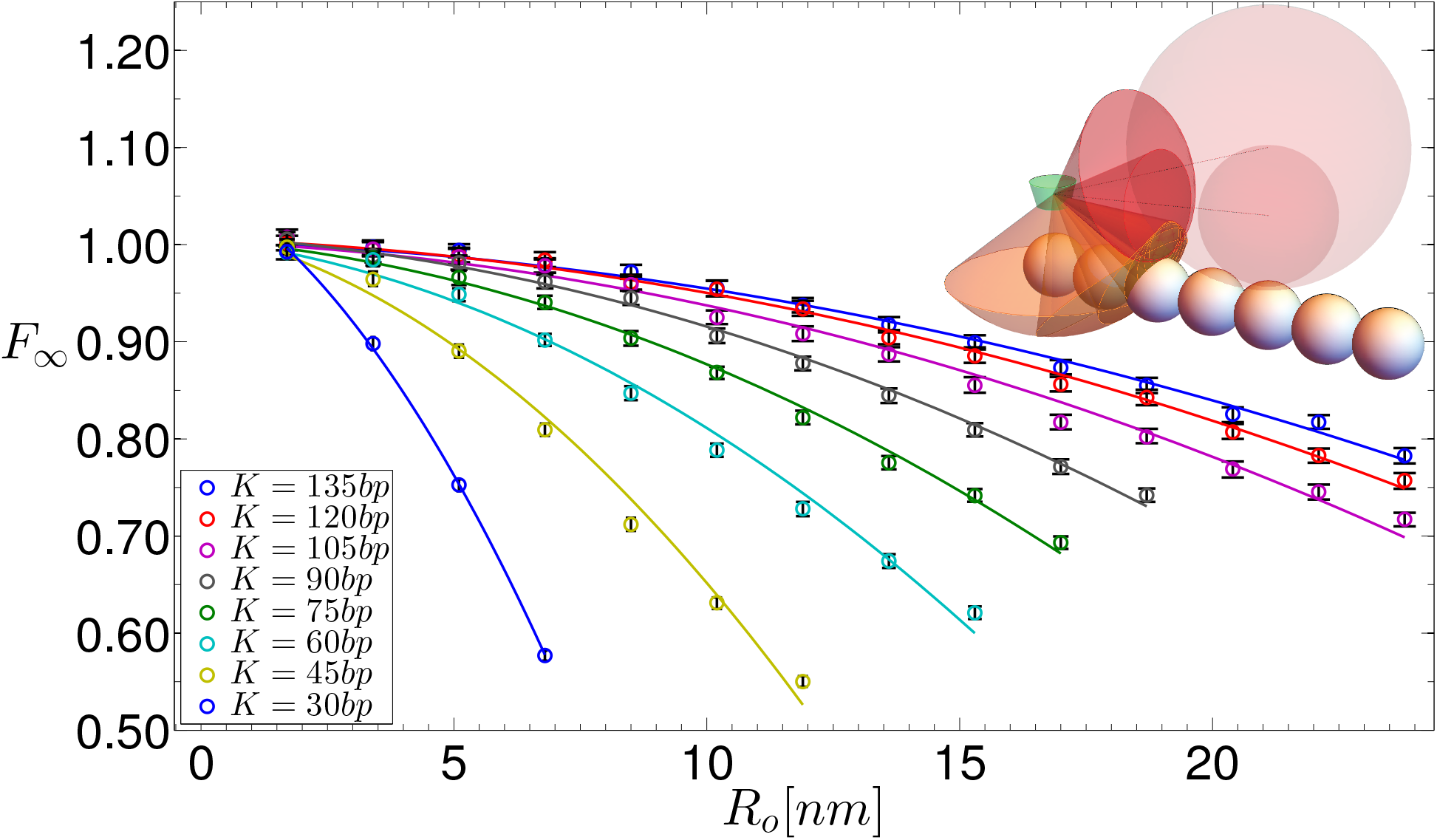
Dependence of *F*_∞_ on protrusion size *R*_o_. Simulation data (circles) are fit (solid curves) by *f_K_* (*R*_o_) (Eq. 15). Data points are the mean of the last 480 points of the simulated *F*. Error bars are ±1.96 times the standard error of these points. Inset illustrates a doubling of R_o_. *δ*r is shown by the green volume. Chain links are shown by white spheres. Protrusions are shown by transparent red spheres. Solid angles subtended by the chain links and protrusion are shown by orange and red cones, respectively. Ω_chain_ is the area on the unit sphere around the center of δr intersecting the orange cones. 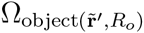 is the area on the unit sphere intersecting the red cone. 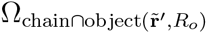 is the area on the unit sphere intersecting both orange and read cones.

In Fig. 4 we plot *F*_∞_ as a function of an on-chain object of radius *R*_o_, for several values of *K*. To compare the results of the numerical simulation to the full eclipsing model (Eq. 12), we first note that when *K* is kept constant, 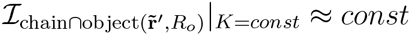, as can be seen from the inset in Fig. 4: the overlap between Ω_chain_ (orange cones) and Ω_object_ (red cones) changes only slightly when the object grows by a factor of two. Furthermore, 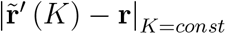 is approximately inde-pendent of *R*_o_ if 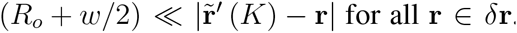. Thus, the dependence of *F*_∞_ on the radius *R*_o_ of an on-chain object can be derived from Eq. (12):

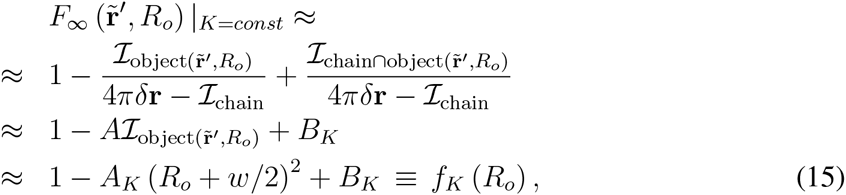

where we approximated 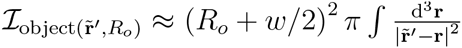 using Eq. (14) and 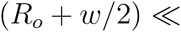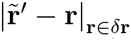. In Fig. 4, we fit the numerical results for different values of *K* with functions of the form *f*_*K*_ (*R*_o_) (Eq. (15). The fits are in excellent agreement 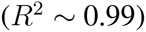 with the numerical data.

**Figure 5:**
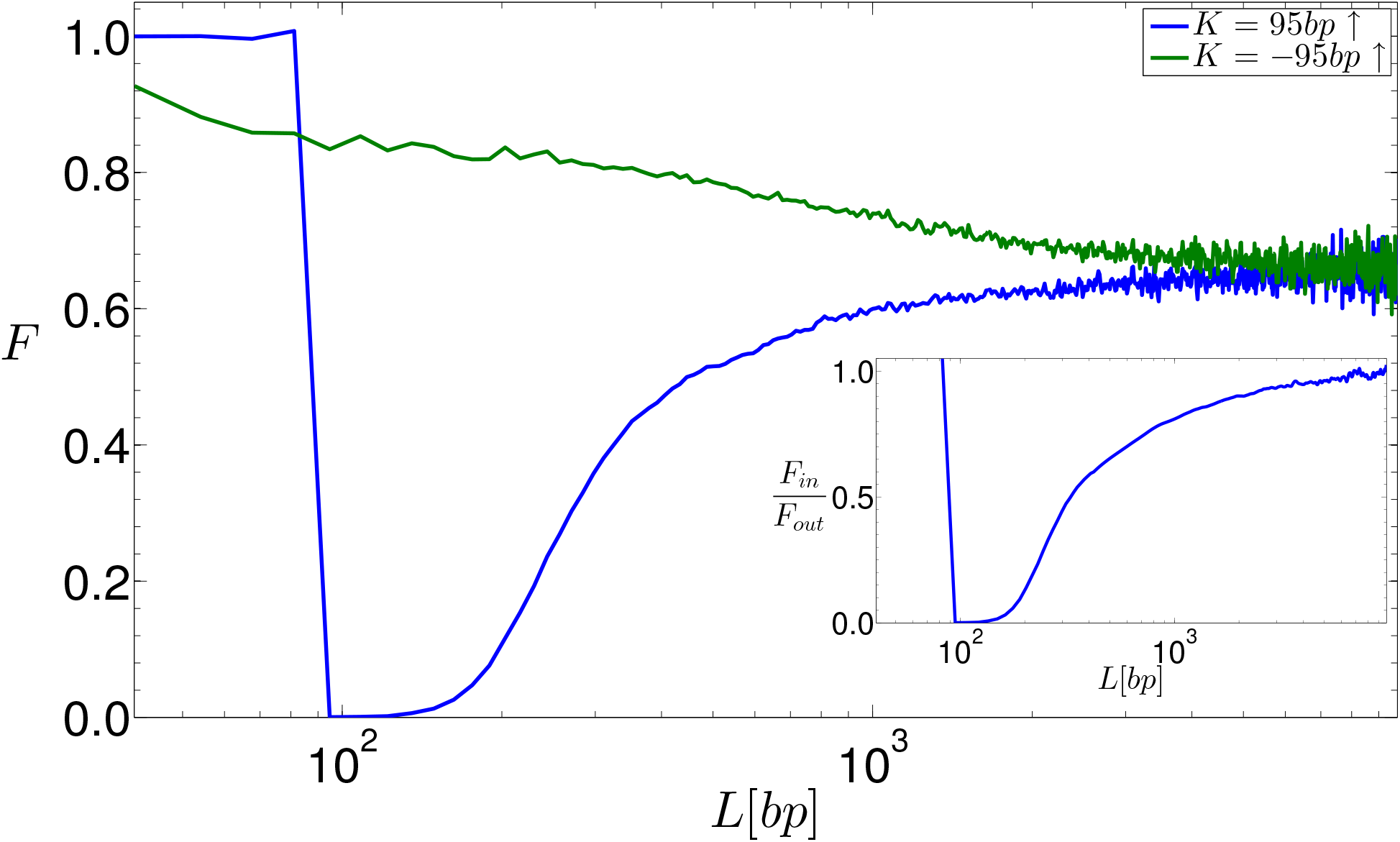
Dependence of *F* (*L*) on location of the protrusion, either inside (*K*) or outside (*K*) the chain segment between link 0 and the terminating link. *R*_o_ = 23 nm and *k* = ±95 bp. Inset: the ratio *F* (*L*,*K*) /*F* (*L*–*K*).

### “Eclipsing” can occur both upstream and downstream of an activator

A salient feature of enhancers is that binding sites for TFs are positioned both upstream and down-stream of the activators. The eclipse model predicts that 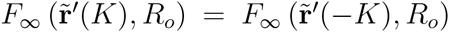, since for long chain lengths the correlation between the positions of both chain termini is completely abolished. Thus, from the perspective of one terminus, looping events can be initiated from any possible direction. To see if our simulation captures this symmetry, we explored a geometry in which the protein-like protrusion was positioned at negative *K* values, corresponding to a location outside of the looping segment delimited by the looping-volume (activator) and distal-terminus (promoter) locations. To do so, we generated an additional chain segment of length Q in the direction opposite to 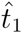, starting from link 0, where *Q* ≫ *K*. We plot the results in Fig. 5. The data show that in the elastic regime, the “outside” geometry (green line, *K* < 0) generates a significantly smaller effect on the probability of looping as compared with the “inside” architecture (blue line, *K* > 0). This lack of symmetry is probably due to the propensity of short loops to form a “tear-drop” shape, thereby reducing the quenching effect of any elements bound outside the looping segment (37). However, for sufficiently large *L*/*b*, *F* (*L*) for both ±*K* enhancer geometries converge to the same value, as predicted by the eclipse model.

### Looping at chain ends vs. looping in the middle of the chain

So far, we modeled an enhancer-promoter region by a chain of discrete semi-flexible links, with the activator and the promoter located at both ends of the chain. However, *in vivo*, the DNA chain extends far beyond the enhancer-promoter region in both directions, and is subject to confinement. We address the first issue in this section, and the issue of DNA confinement in the next section.

We computed the probability of looping for a generic loop of length *L*, located between two internal links of the chain. To do so, we extended the original length of the chain *L* by two flanking segments of 10 Kuhn lengths (10*b*) on both ends of the chain. We assigned the cumulative Rosenbluth weight of the extended chain to the central segment of length *L* containing the enhancer-promoter region. See Extended Materials and Methods, Section 1.3.

We ran a simulation with *K* = 60 bp and *R*_0_ = 11.9 nm (Fig. 6, magenta line) and compared *F* (*L*) to the same case without flanking segments (Fig. 6, blue line). The data show that there is a discrepancy in the long-range looping probability ratio between the two looping models. In particular, the down-regulatory effect of the protrusion is stronger when the enhancer-promoter region is modeled as part of a larger chain. In order to determine the individual contributions of the two flanking segments to the discrepancy, we ran additional simulations with different flanking-segment configurations. We found that only the trailing segment beyond the distal terminus of the looping segment contributes to this discrepancy. The addition of a single chain link of length *w* = 4.6 nm (0.092*b*) after the terminus of the looping segment (Fig. 6, green line) diminishes *F* (*L*), accounting for approximately half of the discrepancy. The addition of only 4 links (0.368*b*) after the terminus of the looping segment (Fig. 6, aqua line) accounts for the entire discrepancy, fully agreeing with the results for the case with 10b flanking segments on both ends of the looping segment. While an addition of a leading segment before the looping volume diminishes the looping probability (data not shown), it does not alter the looping probability ratio *F* (*L*). This can be seen from the comparison between *F* (*L*) for the case with a single link after the terminus of the looping segment (Fig. 6, green line) and the case of a segment of length b before the looping volume and a single chain link after the terminus of the looping segment (Fig. 6, red line).

We believe that these results depend strongly on the looping conditions. For the conditions used here (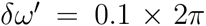, see Fig. 1) the leading flanking segment is inconsequential to the looping probability ratio. However, utilizing a uniform looping volume (*δw*^′^ = 4) would lead to a diminished looping probability ratio as a result of leading segment addition. Similarly, the trailing flanking segment diminishes *F* (*L*) due to the fact that chain configurations that approached the looping volume from a direction colliding with the chain are no longer possible when the chain is extended by a trailing segment. If, however, the looping conditions were such that the chain terminus is required to approach the looping volume from a direction perpendicular to the looping volume cone axis, the trailing segment would become inconsequential to the looping probability ratio.

These results suggest that while there is a discrepancy between the two simulation modes (with flanking regions and without them), it arises from regions that are immediately adjacent to the enhancer-promoter region, and further downstream or upstream segments of the chain do not contribute to the down-regulatory effect.

**Figure 6:**
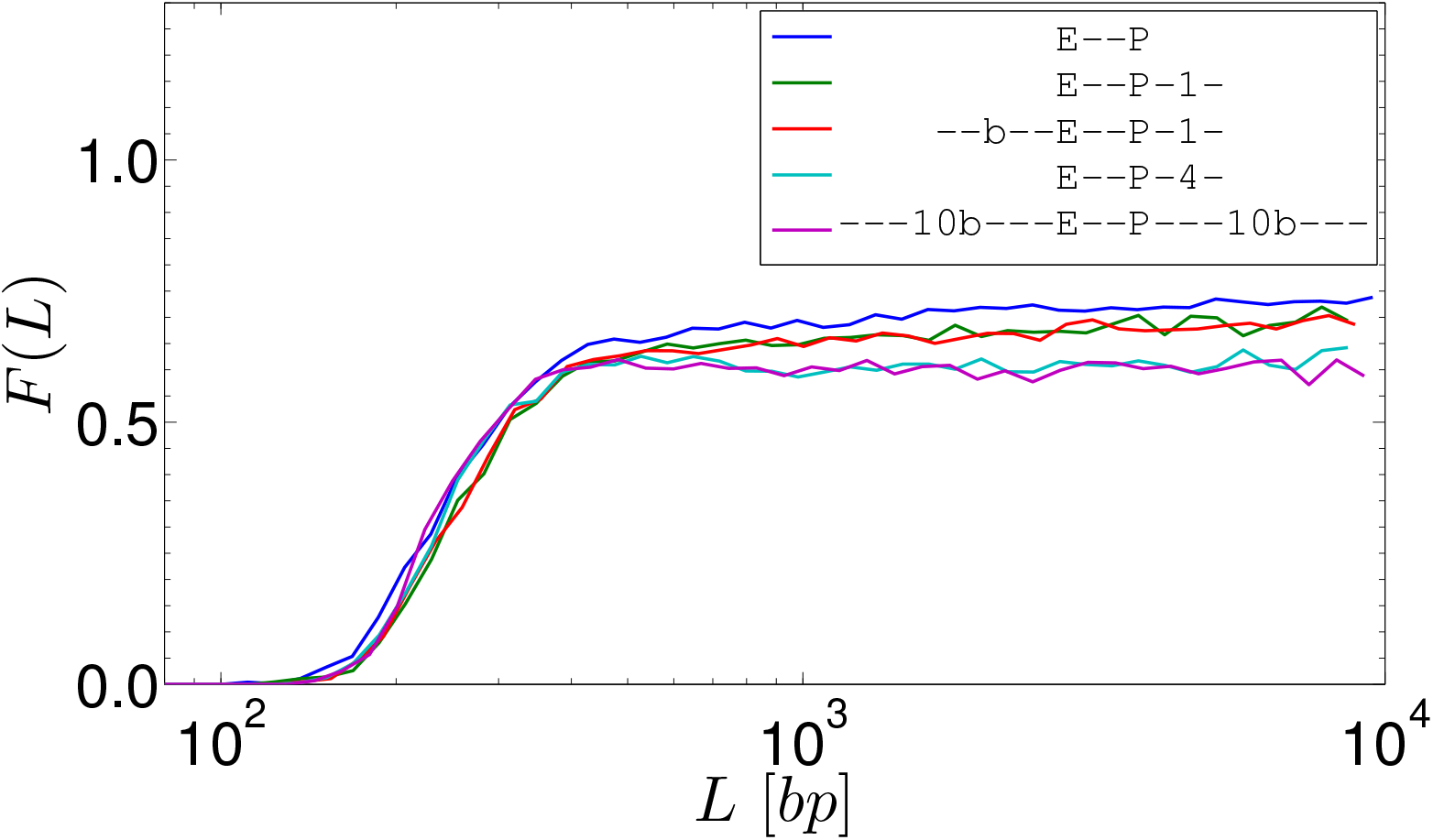
Comparison of *F* (*L*) between enhancer-promoter regions with different-sized flanking segments. All simulations ran with *K* = 60 bp and *R*_0_ = 11.9 nm. Blue: No flanking segments. Green: One link *w* = 4.6 nm (0.092*b*) after the chain terminus. Red: Medium-sized segment of length *b* before the looping volume and one link (0.092*b*) after the chain terminus. Aqua: 4 links (0.368*b*) after the chain terminus. Magenta: Long flanking segments of length 10b on each side of the enhancer-promoter region.

### Looping probability ratio for confined polymer

Chromosomal DNA is fundamentally different from the idealized version assumed in the simulations presented thus far. Cellular DNA is confined, and as a result is compacted into a globular state. This state is characterized by its local nature of fluctuations, as opposed to the coiled state modeled in this work, in which the fluctuations are of the order of the size of the macromolecule itself (40). The nature of the globular state varies between organisms (equilibrated globule in yeast vs. crumpled or fractal globule in eukaryotes (33, 41)) and spans a wide range of volume fractions (35). For lower volume fractions, DNA can be viewed as a semi-dilute polymer solution, in which coils are strongly overlapping but the volume fraction of the polymer in the solution is small. This semi-dilute solution may be pictured as a system of domains (blobs) of characteristic size ξ. Inside each blob, the chain behaves as an isolated macromolecule with excluded volume, and different blobs are statistically independent of one another (8, 40). If the volume fraction is sufficiently low, linearized chromosomal DNA within a blob can explore nearly the entire volume of the blob without interacting with other parts of the chromosome. To simulate this scenario we generated chains confined by spheres of various radii corresponding to a range of realistic blob sizes (35) (for full simulation details, see Extended Materials and Methods).

In Figure 7, we present the normalized looping probability and the looping probability ratio for the range of confining-sphere radii 125-625 nm. Note that the looping probability increases relative to that of the unconstrained polymer when the gyration radius of the chain becomes comparable to the size of the confining sphere (Fig. 7A). However, as can be seen in Fig. 7B, the introduction of a bound protrusion does not influence the looping probability ratio *F* (*L*) up to a chain length of 10 kbp for confining spheres with radii ≥250 nm. For the smallest confining sphere (radius of 125 nm) we were only able to simulate chains of length ≤4.5 kbp (see Supplementary Information Section 3.4), and here too the looping probability ratio remained unaffected. This implies that our results are valid for ~10 kbp of linearized DNA localized within a blob of semidilute chromatin. For additional discussion see Supplementary Information.

We may think of the results of Fig. 7 in a different way. By scaling arguments the simulation also describes a polymer with width of 10 nm (*w* 10=4.8), length of ~6800 nm (*L* × 10/4.8) and persistence length of ~110 nm (*l*_*p*_ × 10=4.8), which is comparable to ~100 kbp of 10 nm chromatin fiber if we assume a density of 15 bp/nm for the fiber (42). The confining sphere forces a specific volume fraction on the confined polymer. The range of confining radii and chain lengths corresponds to polymer volume fractions of up to 0.2%. This interpretation implies that our previous results may be applicable to looping of chromatin fiber at low volume fractions.

### “Eclipsing” effect for *eve* 3/7 stripe

Finally, we applied our model to a real enhancer-promoter system. To do so, we make the following assumptions (based on current experimental evidence). First, when the promoter is active, the enhancer and the promoter region are depleted of nucleosomes for up to 500-1000 bp both upstream and downstream from the center of each regulatory segment (43). This implies that if the enhancer and promoter are sufficiently close, the entire segment can be assumed to be mostly bare dsDNA that is decorated by some DNA binding proteins or protrusions. Second, chromatin in the nucleus is in a globular state. Due to locality of concentration fluctuation in the globular state, we assume that the linearized DNA segment, if not too long, remains trapped in a blob. In order to apply our model to a system with these assumptions, we had to choose an experimentally well-characterized enhancer-promoter system, with an enhancer-promoter separation <10 kbp (consistent with the results presented in Fig. 7B). A likely candidate is the well-characterized *eve* 3/7 enhancer of *D. melanogaster*, which is separated from its promoter by about 4 kbp. In addition, the volume fraction for *D. melanogaster* is estimated between 0.1-1.5% (35). Together, this places the *eve* 3/7 enhancer within the applicability region of our model.

**Figure 7:**
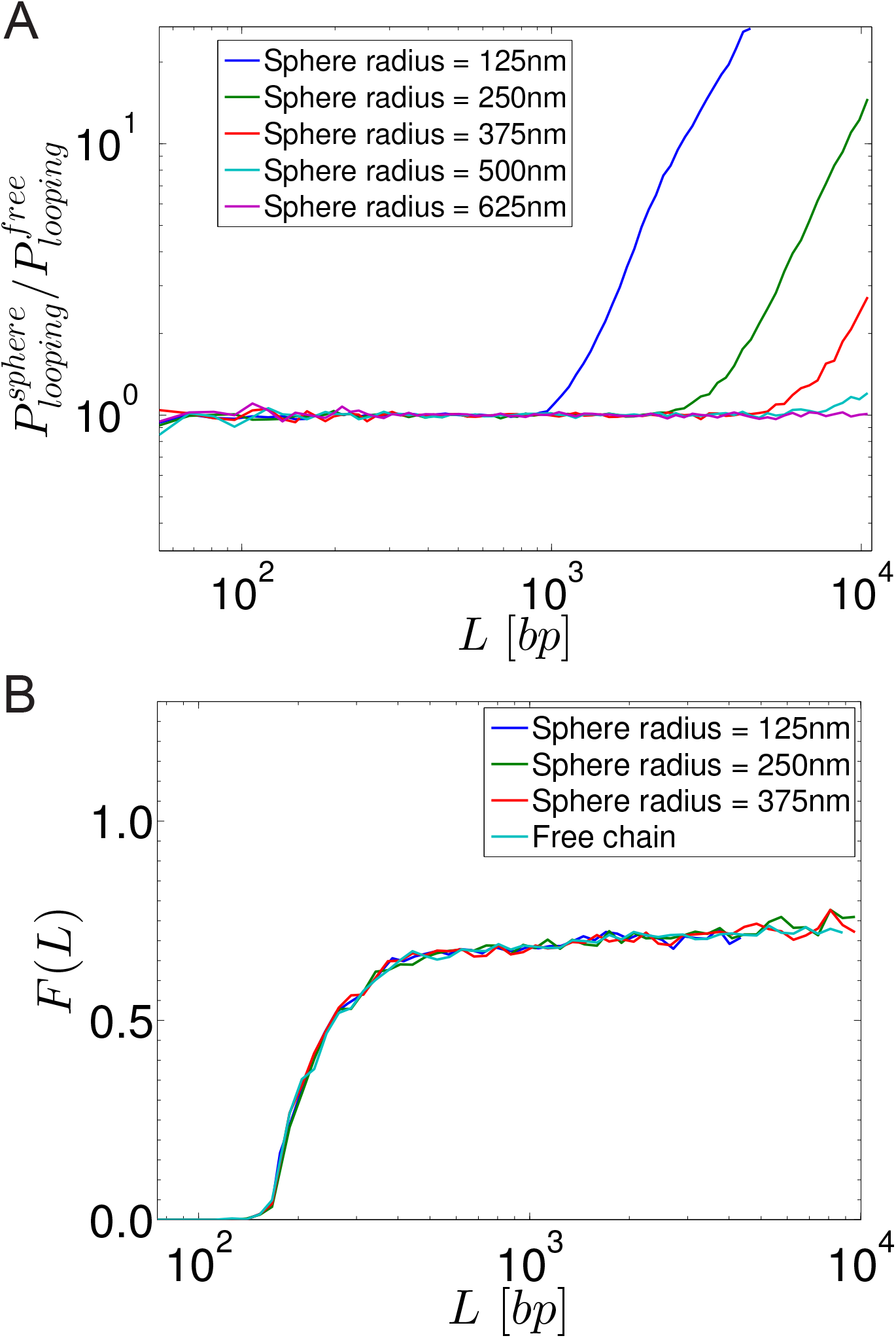
Effect of a confining sphere on the probability of looping. A) Values of 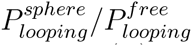 for various sphere sizes. B) Values of *F* (*L*) for three of the smallest spheres compared to *F* (*L*) for the chain without a confining sphere.

We computed our model’s predictions for the active *eve* 3/7 enhancer, i.e. in the early developmental stages (before gastrulation - stage 14), and in the anterior region of the *D. melanogaster* embryo. To check whether the resultant probability of looping could provide a mechanistic explanation for the observed expression pattern, we assumed two possible states: the enhancer is fully occupied by the activators dStat, Zld and the Knirps repressor–co-repressor complexes, and a second state where the enhancer is entirely devoid of bound Knirps repressor complexes. We tested three possible repressor complex sizes: Knirps alone (46 kDa), Knirps bound to CtBP dimers (130 kDa), and a full putative 450 kDa complex, as reported by (23). Since the exact looping conditions of the enhancer looping are unknown, we considered three choices of !^0^ (see Materials and Methods) that represent different extreme cases.

The results, plotted in Fig. 8, show that for all chosen looping conditions, there is a decrease in the probability of enhancer looping when the enhancer is fully bound by Knirps. However, only the largest Knirps–co-repressor complex (containing dCtBP and the HDACs Rpd3 (23) and Sin3 (24)) is large enough to generate a significant reduction in looping.

**Figure 8:**
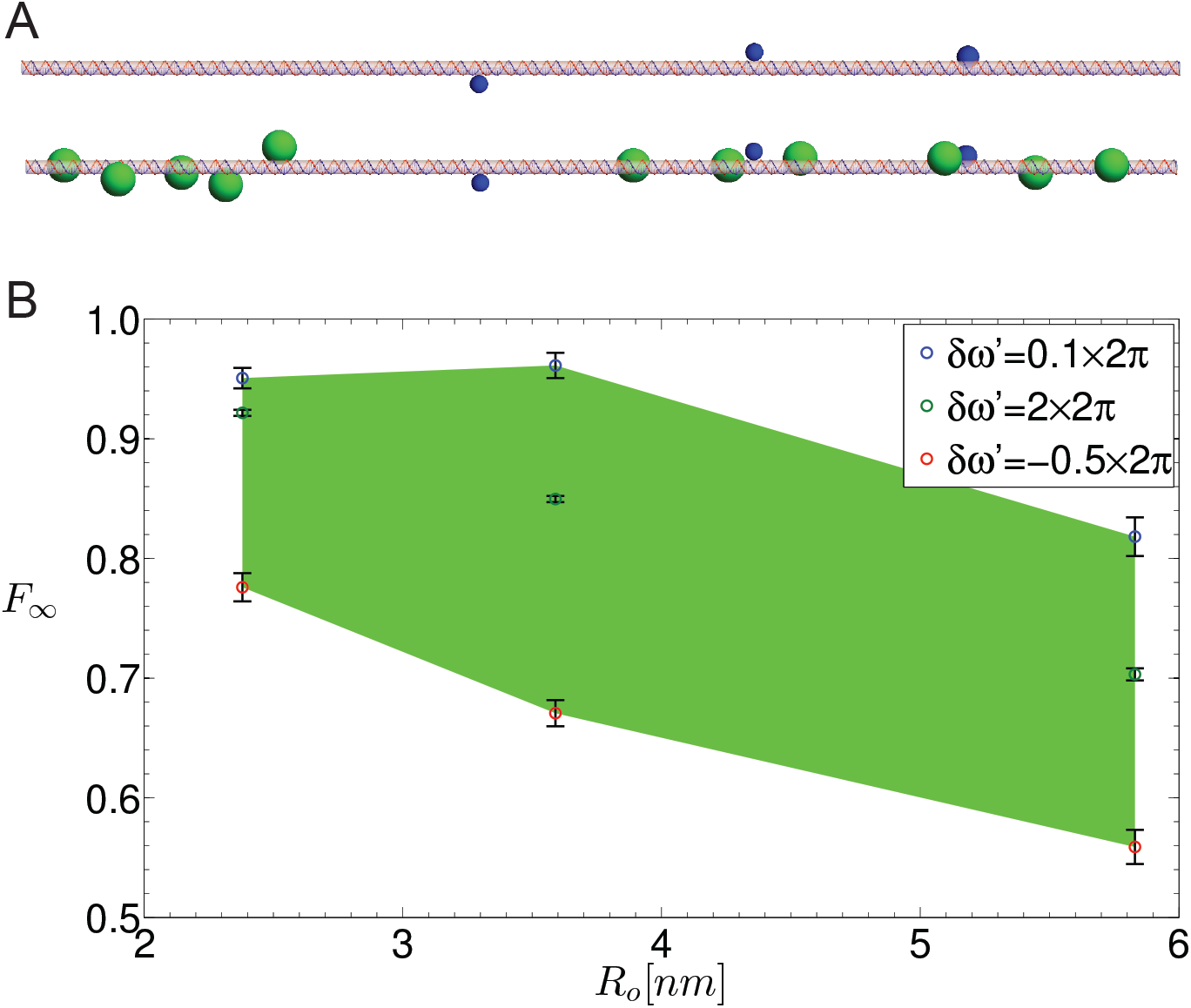
Simulation of the *eve* 3/7 enhancer in *D. melanogaster*. A) Illustration of the *eve* 3/7 enhancer geometry with bound activators (top) vs. with both activators and full putative Knirps+dCtBP+Rpd3+Sin3 complexes (450 kDa) bound (bottom). B) Dependence of *F*_∞_ on protrusion size *R*_o_ and the looping conditions *δ*w^’^. Negative *δ*w^’^ corresponds to anti-collinear orientation of (r_*N*_ r_0_) with 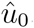. Here *d*_min_ = *R*_chain_ + *R*_activator_ and ε = 3 nm. *F*_∞_ was computed from 150 simulated data points corresponding to DNA chains of lengths 3751-3900 bp. Error bars are ±1.96 times the standard error of these points. The area shaded in green depicts the range of possible *F*_∞_ values, depending on the chosen looping criteria.

## DISCUSSION

We previously established (37) that a bound protein inside a loop can alter the probability of looping in a manner proportional to its size, when the chain length is of the order of the Kuhn length (i.e. < 300 bp). The simulations and theory presented here extend this result to much longer chain lengths. In particular, our model predicts a decrease in the probability of looping that is independent of chain length for long chains in the entropic regime (i.e. *L* ≫ 300 bp), provided that a sufficiently large protrusion oriented in-phase with the looping volume δr is positioned within one Kuhn length of one of the chain termini. We further showed that the reduction in looping probability resembles an eclipse-like phenomenon, where the protrusion blocks the line of sight of one chain terminus from the other.

Our model provides a biophysical mechanism for so-called short-range repression or quenching in enhancers (16–19), which successfully captures many of this phenomenon’s salient features. These include lack of dependence on chain length for sufficiently long chains (*L* ≫ *b*), symmetry with respect to binding-site positioning inside or outside of the looping segment, and the dependence of the regulatory effect on both the size of the bound complex and the distance of the TF from the nearest terminus (~150 bp) for which significant quenching can be generated. To explore the relevance of our model to natural systems, we computed the looping probability ratio for the *D. melanogaster eve* 3/7 enhancer-promoter system in the segment of the embryo where the repressor Knirps is expressed. We found that using realistic repressor-co-repressor-HDAC complex sizes for the simulated protrusions, a significant reduction in looping probability can be obtained, showing that the looping-based mechanism can account for at least some of the quenching generated by Knirps in the context of early fly development. Since there is also a substantial body of evidence supporting HDAC catalysis as a major mechanism for repression (44), we conclude that it is likely that both HDAC activity and looping-related effects combine to generate the quenching observed in *D. melanogaster* and other organisms.

There have been previous attempts to model enhancer regulatory logic for the gap genes of early fly development (45–47). Using a semi-empirical approach, which coupled a thermodynamic model to an empirically-based regulatory scoring function, these works demonstrated that TF occupancy of enhancers can determine the gene-expression outcome. By contrast, our model does not compute the actual expression pattern, but rather the probability of looping for a given occupancy state. For this computation, our model requires only knowledge of the enhancer structure (TF binding sites, protein-complex sizes, and over-all enhancer-promoter distance). This implies that no gene-expression experimental data is needed as an input to predict the down-regulatory effect of bound TFs.

Finally, we consider the range of applicability of our model for natural genomes. While this work focused on bare dsDNA with a protrusion, our results are applicable to any linear polymer as long as its length is significantly larger than its Kuhn length. However, to apply our results to chromosomal DNA, other constraints must be taken into account, some of which we attempted to address in this paper. First, we showed that looping of a linear segment with additional upstream and downstream flanking segments qualitatively resembles looping without flanking segments. Second, chromosomal DNA is subject to confinement, which results in a globular state of the polymer. While there is evidence that active enhancers and their surrounding regions are linear, they are confined by the rest of the chromosome. We attempted to model this constraint using a confining sphere, which allowed us to show that the over-all regulatory effect remains unchanged up to a chain length of ~10 kbp for sphere sizes corresponding to realistic blob sizes. Third, DNA is typically a mixture of bare and chromatinized DNA. While we did not address the issue of “mixed” DNA directly, we do know that chromatinized DNA is more densely packed than bare DNA. Thus, the ~10 kbp looping results for bare DNA correspond to much longer lengths (up to ~100 kbp) of “mixed” DNA. We can conclude that the ~10 kbp range for bare DNA addressed by our simulations is a lower bound on the actual range of validity of our model for “mixed” or chromatinized DNA. Our work indicates that we should consider three ranges for the physics of looping of chromatinized DNA. For short ranges which we studied previously (14, 37), looping is dominated by elastic energy. As shown in this work, there is an intermediate entropic looping range of up to ~10 kbp of bare DNA (or ~100 kbp of “mixed” DNA). Recent studies indicate that there may be an additional long-range regime in which interactions between neighboring regions of the globular DNA must be taken into account (33, 48, 49).

## AUTHOR CONTRIBUTIONS

YP devised and wrote the simulations. YP, SG, and RA analyzed the data and wrote the manuscript.

## ACKNOWLEDGEMENTS

This project was funded by the European Union’s Horizon 2020 Research And Innovation Programme under grant agreement 664918–-MRG-GRammar, by the Israel Science Foundation through grant 1677/12, by the I-CORE Program of the Planning and Budgeting Committee, and by the Israel Science Foundation (grant 152/11). Y.P. acknowledges support provided by the Russell Berrie Nanotechnology Institute, Technion. The authors declare no conflicts of interest.

## SUPPLEMENTARY MATERIAL

An online supplement to this article can be found by visiting BJ Online at http://www.biophysj.org.

